# Genome-wide analyses of PAM-relaxed Cas9 genome editors reveal substantial off-target effects by ABE8e in rice

**DOI:** 10.1101/2022.02.09.479813

**Authors:** Yuechao Wu, Qiurong Ren, Zhaohui Zhong, Guanqing Liu, Yangshuo Han, Yu Bao, Li Liu, Shuyue Xiang, Shuo Liu, Xu Tang, Jianping Zhou, Xuelian Zheng, Simon Sretenovic, Tao Zhang, Yiping Qi, Yong Zhang

## Abstract

PAM-relaxed Cas9 nucleases, cytosine base editors and adenine base editors are promising tools for precise genome editing in plants. However, their genome-wide off-target effects are largely undetermined. Here, we conduct whole-genome sequencing (WGS) analyses of transgenic plants edited by xCas9, Cas9-NGv1, Cas9-NG, SpRY, nCas9-NG-PmCDA1, nSpRY-PmCDA1 and nSpRY-ABE8e in rice. Our results reveal different guide RNA (gRNA)-dependent off-target effects with different editors. *De novo* generated new gRNAs by SpRY editors lead to additional but not substantial off-target mutations. Strikingly, ABE8e results in ~500 genome-wide A-to-G off-target mutations at TA motif sites per transgenic plant. The preference of the TA motif by ABE8e is also observed at the target sites. Finally, we investigate the timeline and mechanism of somaclonal variation due to tissue culture, which chiefly contributes to the background mutations. This study provides a comprehensive understanding on the scales and mechanisms of off-target and background mutations during PAM-relaxed genome editing in plants.

---

CRISPR-Cas9 genome editing tools have greatly revolutionized plant genetics and breeding. *Streptococcus pyogenes* Cas9 (SpCas9) is the predominant Cas9 widely used, partly due to its high genome editing efficiency and simple NGG protospacer adjacent motif (PAM) requirement[1–3]. To broaden the targeting scope, many SpCas9 variants have been engineered, including xCas9 (recognizing NG, GAA and GAT PAMs)[4], SpCas9-NGv1 and SpCas9-NG (recognizing NG PAM)[5], and PAM-less SpRY[6]. These PAM-relaxed Cas9 nucleases have been widely adopted for genome editing in plants[7, 8]. However, their relaxed PAM requirements could make them prone to guide RNA (gRNA)-dependent off-targeting, which awaits a comprehensive investigation in plants.

The development of cytosine base editors (CBEs) and adenine base editors (ABEs) further expanded the genome editing toolbox[9], enabling precise base changes in plants[10]. Cytidine deaminases and adenosine deaminases used in CBEs and ABEs could potentially catalyze deamination reactions nonspecifically in the genomes, causing gRNA-independent off-target effects. For example, whole-genome sequencing (WGS) revealed such off-target effects for rAPOBEC1-based CBEs in rice[11, 12] and mouse[13]. CBEs engineered with different cytidine deaminases showed less off-target effects in human cells[14, 15] and in plants[12, 16]. ABE8e, a highly processive ABE[17], catalyzes highly efficient A-to-G base transitions in human cells[18] and in plants[19–23]. Although elevated A-to-I conversions were reported in the transcriptomes of ABE8e-treated human cells[18], it is unknown whether or to what extent gRNA-independent off-target mutations in plants would be generated by ABE8e.

Merger of PAM-relaxed Cas9 variants and highly efficient cytidine/adenosine deaminases opens the door for highly flexible base editing in plants[10]. CBEs based on xCas9 were reported in plants to edit NGN PAM sites, albeit with very low efficiency[24–27]. SpCas9-NGv1 and SpCas9-NG based CBEs were tested in different plant species[24–26, 28, 29], generally outperforming xCas9 based CBEs at relaxed PAM sites[10]. SpRY CBEs were demonstrated to edit NRN PAMs better than NYN PAMs in rice[19–21, 30]. Similarly, ABEs were demonstrated in plants with SpCas9-NGv1[31] or SpCas9-NG [24, 26, 32] and SpRY[19–21, 30, 33]. Despite the wide demonstration of these PAM-relaxed CBEs and ABEs in plants, their potential genome-wide off-target effects have not been reported. To fill this critical knowledge gap, we comprehensively assessed gRNA-dependent and -independent off-target effects of these PAM-relaxed nucleases and base editors using WGS in rice. We also investigated the generation of somaclonal variation in the context of genome editing.

## Results

### The experimental pipeline for studying off-target effects of PAM-relaxed genome editing in rice by whole-genome sequencing

Our previous study revealed that xCas9 largely retained the NGG PAM requirement of SpCas9 with improved editing specificity[25]. To simply validate this observation, we included an xCas9 construct for editing an NGG PAM site with OsDEP1-gR02-GGG. Although SpCas9-NGv1 and SpCas9-NG both recognize NGN PAMs[5, 29, 31], SpCas9-NG has higher editing efficiency than SpCas9-NGv1[5, 25]. It is intriguing to compare SpCas9-NGv1 and SpCas9-NG variants for their off-target effects and, hence, we targeted two independent sites OsDEP1-gR01-GGT and OsDEP1-gR02-CGC with both variants. Since genome-integrated T-DNAs are prone for self-editing by SpRY and its derived base editors[19], we wanted to investigate the scale of off-target mutagenesis due to such *de novo* generated gRNAs by SpRY at four different target sites (OsDEP1-gR01-CGC, OsDEP1-gR04-CGC, OsPDS-gR01-TCA and OsPDS-gR03-TAA). For off-target analysis of PAM-relaxed CBEs, we focused on SpCas9-NG and SpRY with a highly efficient and specific PmCDA1 cytidine deaminase[12]. This allows us to focus our analysis on gRNA-dependent off-target effects of nSpCas9-NG-PmCDA1 and nSpRY-PmCDA1 with each editing two target sites (OsDEP1-gR01-TGT and OsDEP1-gR02-CGC for nSpCas9-NG-PmCDA1; OsALS-gR21-GCA and OsALS-gR22-AGC for nSpRY-PmCDA1). By contrast, off-target effects of the highly efficient adenosine deaminase, ABE8e, are largely unknown. Using nSpRY-ABE8e to edit two independent sites (OsPDS-gR01-TGG and OsPDS-gR04-TAA), we hoped to reveal both gRNA-dependent and -independent off-target effects by this highly efficient PAM-less ABE.

These constructs, along with their no corresponding gRNA controls (**Supplementary Table 1**), were used to generate transformed rice plants through *Agrobacterium* mediated transformation. Genome editing frequencies were calculated for most constructs including PAM-relaxed Cas9 nucleases (SpCas9-NGv1, SpCas9-NG and SpRY) (**Fig. 1a**), and CBEs (nSpCas9-NG-PmCDA1 and nSpRY-PmCDA1) (**Fig. 1b**), and nSpRY-ABE8e (**Fig. 1c**). As expected, SpCas9-NG showed higher editing efficiency than SpCas9-NGv1 (**Fig. 1a**). Different numbers (one to four) of genome edited T_0_ lines from different constructs and regenerated T_0_ lines from the no corresponding gRNA constructs were chosen for WGS control samples (**Fig. 1d and Supplementary Table 1**). The resulting sequencing data showed >50X sequencing depth, >99% mapping ratio, and >97% genome coverage for all 58 samples (**Supplementary Table 2**), which were processed according to a rigid bioinformatics pipeline to call out single nucleotide variations (SNVs) and insertions or deletions (INDELs) for further comparisons and analyses (**Fig. 1e**)[12, 34]. We analyzed the three T_0_ lines edited by xCas9 at OsDEP1-gR02-GGG site and did not find gRNA-dependent off-target mutations (**Supplementary Table 3**), which is consistent with its high targeting specificity reported in human cells[4] and in rice[25].

**Figure 1.**
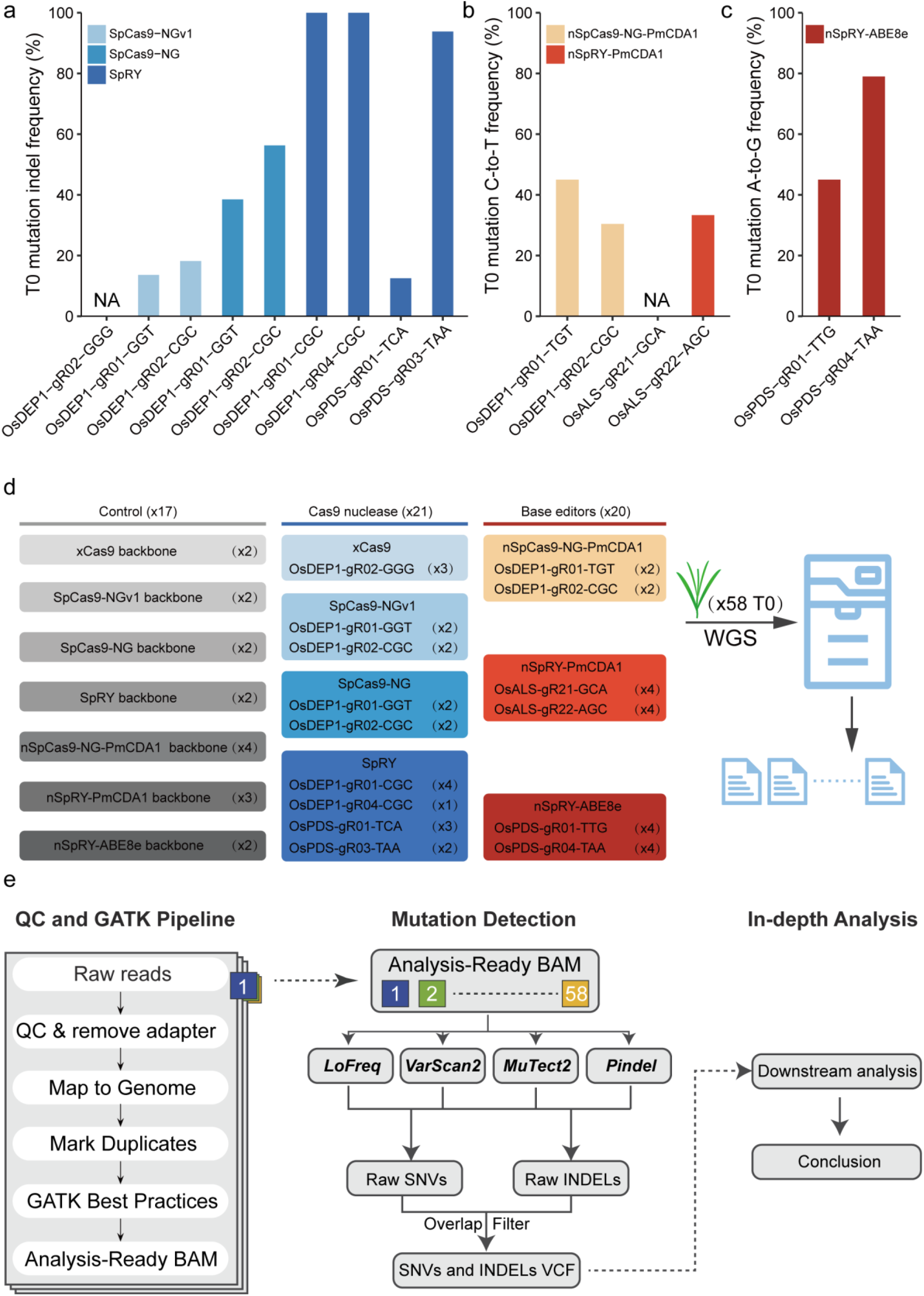
Assessment of PAM-less genome editing in rice by whole-genome sequencing. **a-c**, genome editing frequencies in T_0_ lines by PAM-relaxed Cas9-NGv1, Cas9-NG and SpRY (**a**), by PAM-relaxed cytosine base editors based on nCas9-NG and nSpRY (**b**), and by PAM-less nSpRY-ABE8e adenine base editor (**c**). **d**, Summary of plants used for whole-genome sequencing. **e**, The bioinformatic pipeline for analysis of whole-genome sequencing (WGS) data. NA, editing frequency in T_0_ lines was not scored for the constructs xCas9-OsDEP1-gR02-GGG and nSpRY-PmCDA1-OsALS-gR21-GCA.

### Comparison of SpCas9-NGv1, SpCas9-NG and nSpCas9-NG-PmCDA1 reveals differential gRNA-dependent off-target effects dictated by nuclease activity and editor types

We next compared SpCas9-NGv1, SpCas9-NG and nSpCas9-NG-PmCDA1 at editing NGN PAM sites. At OsDEP1-gR02-CGC site, WGS discovered six off-target sites that were edited by SpCas9-NGv1, five out of six being shared among two T_0_ lines **(Fig. 2a)**. All these six off-target sites contain NGN PAMs and no more than 1 mismatch mutation in the 3-20 nt region of the protospacers, suggesting high likelihood of off-target editing. The resulting off-target mutations are small deletions and 1-bp insertions around Cas9 cleavage site, 3 bp upstream of the PAM (**Fig. 2a**), which are hallmarks of Cas9 editing outcomes. A total of 11 off-target sites with NGN PAMs were discovered among the two T_0_ lines edited by SpCas9-NG, including the four identified with SpCas9-NGv1 (**Fig. 2b**). Only one off-target mutation was shared by the two T_0_ lines (**Fig. 2b**). Many of the newly discovered off-target sites with SpCas9-NG contain two or more mismatches to the protospacer (**Fig. 2b**), which is consistent with increased nuclease activity of SpCas9-NG over SpCas9-NGv1[5, 25]. Six off-target sites were identified in the two T_0_ lines edited by nSpCas9-NG-PmCDA1, with three different off-target sites in each line (**Fig. 2c**). Unlike SpCas9-NGv1 and SpCas9-NG that shared four off-target sites, the six off-target sites identified with nSpCas9-NG-PmCDA1 are all different from those identified with the nucleases (**Fig. 2d**), suggesting gRNA-dependent off-target mutations by Cas9 nucleases and base editors follow different mechanisms. Of note, four of the six off-target sites carry deletions spreading across the protospacer (**Fig. 2c**), supporting the off-target mutations were caused by cytidine deaminase activity and base excision repair. Interestingly, none of the T_0_ lines analyzed here showed evidence of T-DNA self-editing. This could be explained by the fact that the GTT PAM in the gRNA scaffold is not an optimal PAM for SpCas9-NGv1, SpCas9-NG and nSpCas9-NG-PmCDA1, although self-editing by SpCas9-NG was previously reported in rice[35].

**Figure 2.**
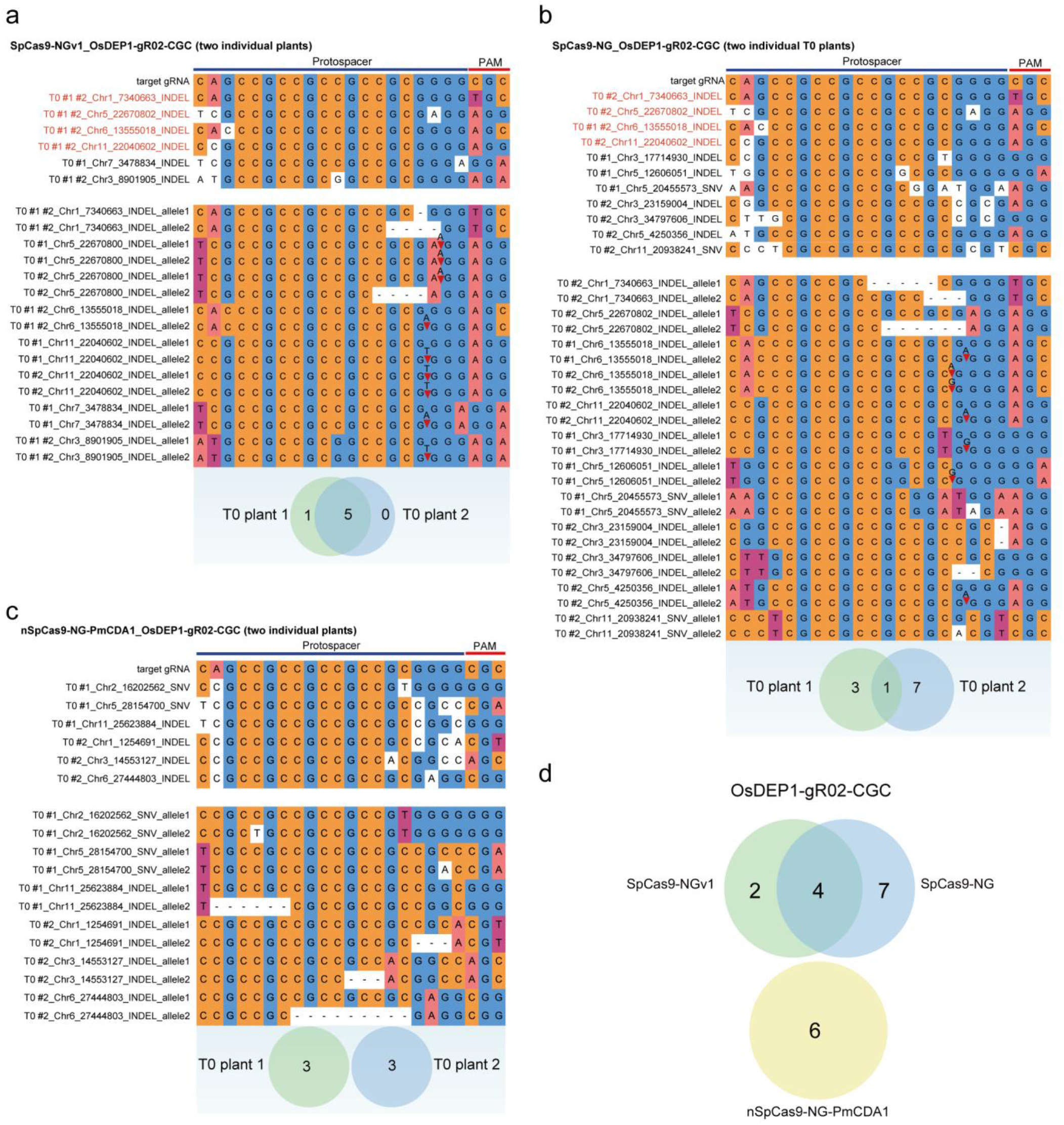
Different sequence preference of gRNA-dependent potential off-target editing by Cas9-NG nucleases and cytosine base editors. **a-c**, gRNA-dependent off-target mutations in edited T_0_ lines at the OsDEP1-gR02-CGC site by SpCas9-NGv1 (**a**), SpCas9-NG (**b**), and nSpCas9-PmCDA1 (**c**). Off-target sites that were shared between SpCas9-NGv1 and SpCas9-NG are marked in red. Top panel, sequence comparison of target gRNA and potential off-target sites. Middle panel, the genotype of the off-target sites. Bottom panel, the number of potential off-target sites in two T_0_ plants. **d**, Venn diagram depicting many shared off-target sites induced by the OsDEP1-gR02-CGC gRNA in SpCas9-NGv1 and SpCas9-NG, while not in nCas9-NG-PmCDA1.

### Comparison of SpRY and nSpRY-ABE8e reveals gRNA-dependent off-target mutations by *de novo* generated gRNAs

To investigate gRNA-dependent off-target effects of SpRY editors, we first investigated the gRNA-dependent off-target effects by SpRY derived base editors. The results showed that no gRNA-dependent off-targeting was found in the edited T_0_ lines by nSpRY-PmCDA1 (**Supplementary Table 3**). However, 18 and 5 potential off-target sites with up to 5 mismatches were edited by SpRY and SpRY-ABE8e, respectively (**Supplementary Table 3**). Among these edited off-target sites, 21 out of 23 contain no more than 3 mismatch mutations in the 3-20 nt region of the protospacers (**Supplementary Fig. 1a-b and Supplementary Fig. 2**). Thus, the off-target effect of SpRY could be minimized by improving the specificity of protospacers.

We next focused our analysis on *de novo* generated gRNAs due to T-DNA self-editing, a common phenomenon caused by the PAM-less nature of SpRY[19]. Ten lines were analyzed at four target sites (**Fig. 1a and 1b**). New gRNAs were generated at all four target sites among eight T_0_ lines (**Fig. 3a and Supplementary Fig. 3**). Based on these new protospacers, we identified potential off-target sites with 0-5 nucleotide mismatches using Cas-OFFinder[36]. However, only two new gRNAs resulted in off-target mutations at these predicted off-target sites (**Fig. 3a**). At OsDEP1-gR01-CGC site, one new gRNA appeared to cause one SNV mutation at a target site with multiple nucleotide mismatches (**Fig. 3b**). Similarly, at OsDEP1-gR04-CGC site, one new gRNA seemed to generate either SNV or INDEL mutations at five off-target sites (**Fig. 3c**). These off-target sites showed significant difference to the protospacer of the original target gRNA (**Fig. 3c**), suggesting that the mutations at these sites were unlikely to be caused by the original gRNA, rather more likely to be created by the new gRNA. Given that detected mutations at these off-target sites are located upstream relative to the Cas9 cleavage site (**Fig. 3b and 3c**), it is possible that some of these mutations might not be caused by gRNA-dependent SpRY editing.

**Figure 3.**
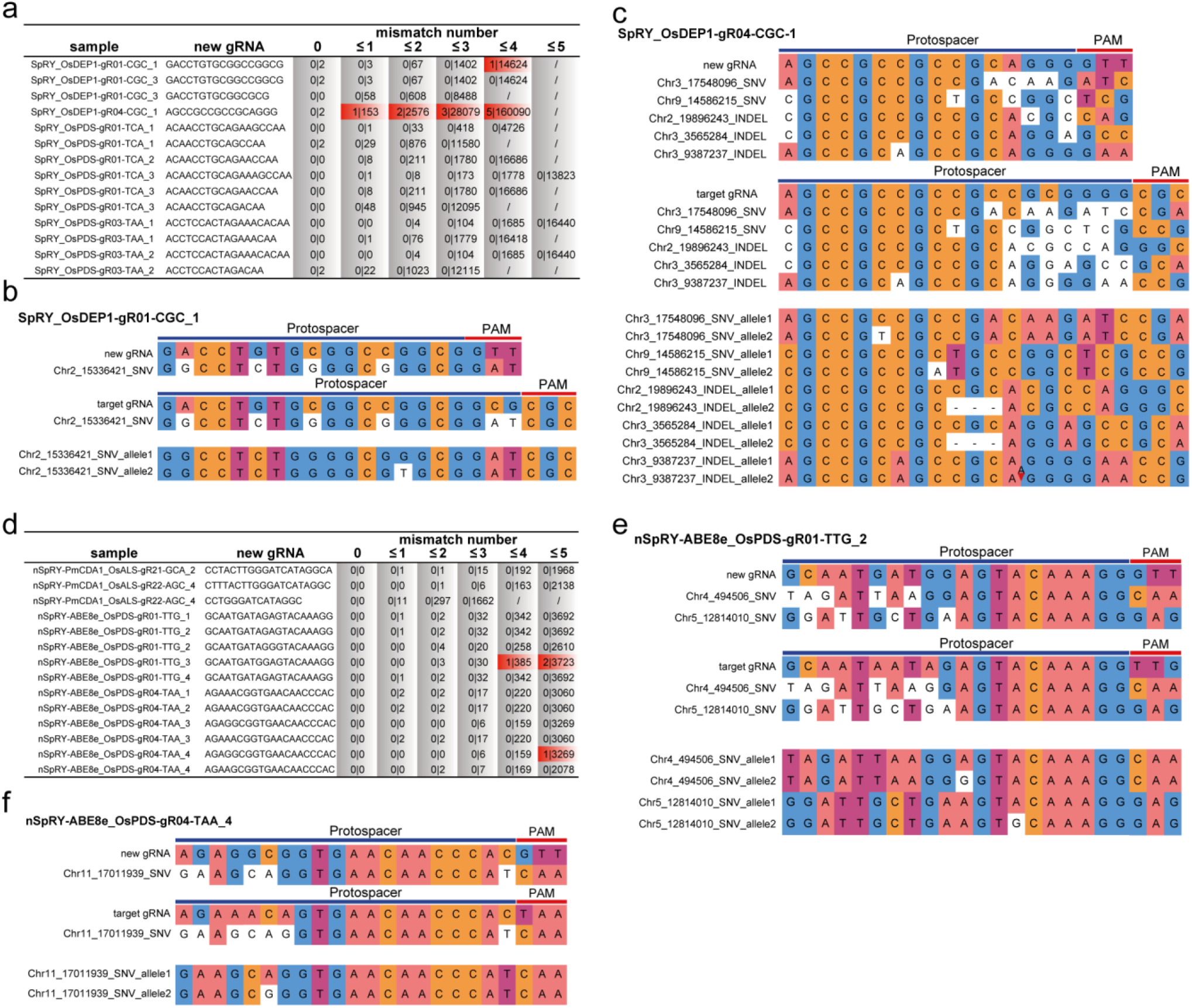
Genome-wide landscape of gRNA-dependent off-target mutations by de novo generated new sgRNAs by SpRY editors. **a, d**, Off-target analysis for *de novo* generated new gRNAs due to on-target editing by SpRY nuclease, nSpRY-PmCDA1 and nSpRY-ABE8e. The number of off-target sites overlapping identified mutation (SNVs+INDELs) versus the number of all potential off-target sites that predicted by Cas-OFFinder. **b-c**, gRNA-dependent off-target mutations in T_0_ lines by *de novo* generated new gRNAs by SpRY at the OsDEP1-gR01-CGC site (**b**) and the OsDEP1-gR04-CGC-1 site (**c**). Top panel, sequence comparison of new gRNA and potential off-target sites. Middle panel, sequence comparison of target gRNA and potential off-target sites. Bottom panel, the genotype of the off-target sites. **e-f**, gRNA-dependent off-target mutations by *de novo* generated new gRNAs by nSpRY-ABE8e at the OsPDS-gR01-TTG-2 site (**e**) and OsPDS-gR04-TAA-4 site (**f**).

We also investigated self-editing related off-target effects of SpRY based CBE and ABE. For nSpRY-PmCDA1, T-DNA self-editing of the OsALS-gR21-GCA construct and the OsALS-gR22-AGC construct was detected in one out of two T_0_ lines each (**Supplementary Fig. 4**), generating one and two new gRNAs, respectively (**Fig. 3d**). For all three new gRNAs, WGS did not detect off-target mutations at the off-target sites predicted by Cas-OFFinder (**Fig. 3d**). For nSpRY-ABE8e, T-DNA self-editing was detected in most T_0_ lines for the OsPDS-gR01-TTG and the OsPDS-gR04-TAA constructs (**Fig. 3d and Supplementary Fig. 5**). Interestingly, in both cases, no off-target mutations were detected at Cas-OFFinder-predicted off-target sites with three or fewer nucleotide mismatches (**Fig. 3d**). However, for nSpRY-ABE_OsPDS-gR01-TTG, mutations were detected in line 2 at two predicted off-target sites with four and five nucleotide mismatches to the protospacer of the new gRNA and with six nucleotide mismatches to the protospacer of the original target gRNA (**Fig. 3e**). Similarly, one off-target mutation was detected for nSpRY-ABE_OsPDS-gR04-TAA in line 4, where the off-target site showed two fewer mismatches (five vs. seven) to the protospacer of the new gRNA than the original target gRNA (**Fig. 3f**). All three off-target events are A-to-G conversions at target sites with NRN PAMs (**Fig. 3e and 3f)**, consistent with high purity base conversion by ABE8e[18] and SpRY PAM preference of NRN PAMs over NYN PAMs[6]. Together, these data suggest that very few gRNA-dependent off-target mutations were induced by PAM-relaxed SpRY base editors.

### Comparison of PAM-relaxed nucleases and base editors reveals gRNA-independent genome-wide off-target A-to-G mutations by ABE8e

We next pursued our analyses to reveal any off-target effects of these PAM-relaxed editors that are independent of gRNAs. For xCas9, SpCas9-NGv1, SpCas9-NG, SpRY and nSpRY-PmCDA1 constructs, both genome-edited plants and control plants shared similar numbers of SNVs (ranging from 86 to 322, on average 187), INDELs (ranging from 48 to 108, on average 75) (**Fig. 4a and Supplementary Fig. 6)** and frequencies of deletions for different sizes (**Supplementary Fig. 7**). These mutations appeared to be present in all genomic regions across the genome (**Fig. 4b and Supplementary Fig. 8**). Importantly, the numbers of SNVs and INDELs observed are in the same range as those observed in other groups and our previous studies[11, 12, 16, 34], supporting these mutations were somaclonal variation due to tissue culture. Strikingly, both genome-edited plants and control plants expressing nSpRY-ABE8e showed many more SNVs, averaging 700 per plant (**Fig. 4a**) and being present in all genomic regions (**Fig. 4b**). By contrast, nSpRY-ABE8e expressing plants showed similar numbers of INDELs (on average 77) to other plant groups (**Supplementary Fig. 6**). A close analysis showed the excessive amount of SNVs in nSpRY-ABE8e expressing plants are A-to-G mutations, and the high enrichment of A-to-G mutations and decreased fractions of other nucleotide substitutions were only observed with plants expressing nSpRY-ABE8e (**Fig. 4c**). These A-to-G mutations were randomly spread across all 12 chromosomes of rice genome (**Fig. 4d**). About 95% of these A-to-G mutations belong to the category of 25%-75% allele frequencies (**Supplementary Fig. 9**), suggesting these are largely germline transmittable mutations. Our results hence demonstrated genome-wide gRNA independent A-to-G off-target mutagenesis in rice by the highly processive ABE8e.

**Figure 4.**
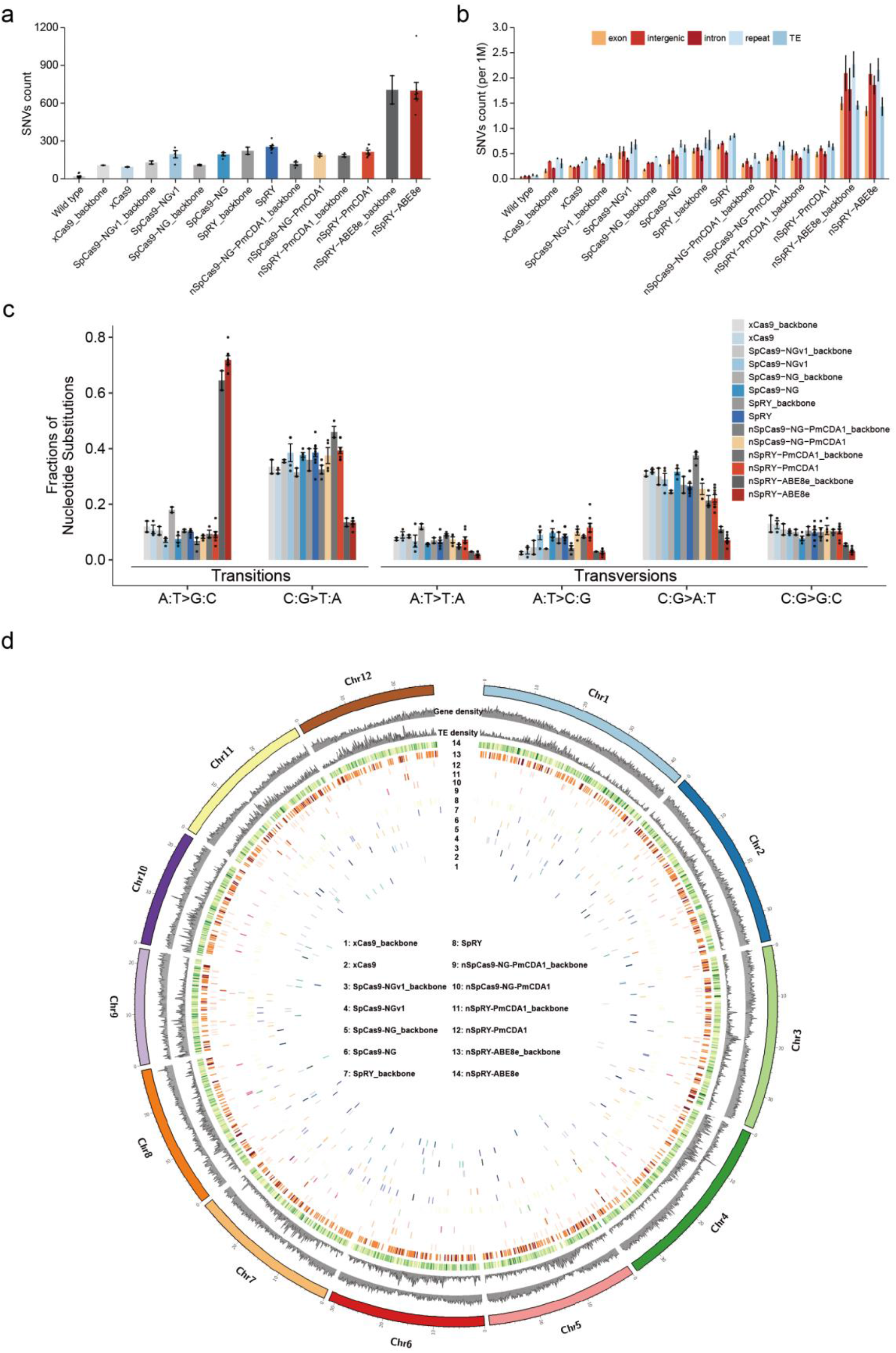
Genome-wide sgRNA-independent off-target effects by PAM-relaxed nucleases, cytosine base editors, and adenine base editors. **a**, Number of single nucleotide variation (SNV) mutations in all sequenced samples. **b**, Average number of SNV mutations in per 1 Mbp genomic region. **c**, Fractions of different nucleotide substitutions in different samples. **d**, Genome-wide distribution of A-to-G SNVs in all sequenced samples. **a-c**, Error bars represent s.e.m. and dots represent individual plants.

### ABE8e favors TA motif sites for both off-target and on-target editing

To further study the off-target effects by ABE8e, we analyzed all the A-to-G off-target editing sites in 10 T_0_ lines. The results showed unambiguously that ABE8e favors conversion of A to G in TA motifs on either Watson strand (**Fig. 5a**) or Crick strand (**Supplementary Fig. 10**). We reasoned that such a preference of editing TA motifs by ABE8e could also be reflected at on-target sites. To this end, we tested nCas9-ABE8e at editing an NGG PAM site in rice protoplasts and the data showed A-to-G conversions at both A_4_ and A_12_ (**Fig. 5b**), with both positions being at the edge of the editing window known for ABE8e[18]. The editing frequency at A_12_ proceeded by a ‘T’ is significantly higher than A_4_ proceeded by a ‘G’ (**Fig. 5b**), supporting that ABE8e also favors TA motifs for on-target editing. We then analyzed all 11 edited alleles in T_0_ lines by nSpRY-ABE8e_OsPDS-gR01-TTG (**Fig. 5c**) and found A_6_ proceeded by a ‘T’ was edited at much higher frequency than A7 proceeded by an ‘A’ (**Fig. 5d**), although both A_6_ and A7 are within the ABE8e editing window. Furthermore, we analyzed the gRNA-dependent off-target editing outcomes discovered at four off-target sites by the same construct (**Fig. 5e**). A-to-G conversions were only found at TA sites, not at AA, CA, and GA sites (**Fig. 5f**). Taken together, these analyses indicate that ABE8e has a strong preference of the TA motif for both off-target and on-target editing.

**Figure 5.**
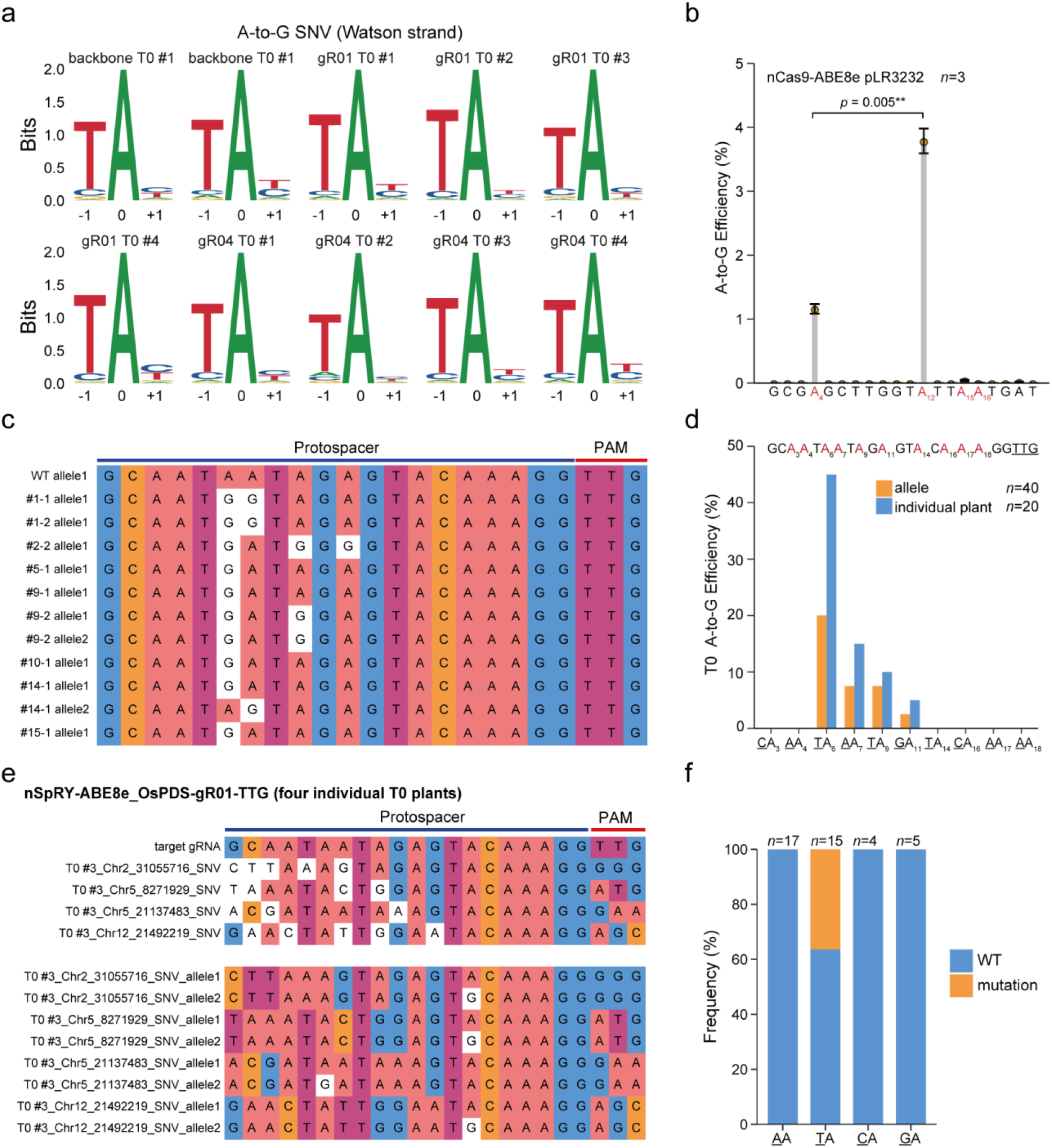
ABE8e favors A-to-G conversion at TA motifs at both off-target and on-target sites. **a**, Preference of a TA motif by ABE8e at gRNA-independent off-target A-to-G base editing in Watson strand, 0 indicates the A-to-G SNV position. **b**, Base editing frequencies at different protospacer positions by ABE8e at a target site in rice protoplasts, *n* represents biological replicates. Data reanalyzed from ref[19]. Error bars represent s.e.m. *p*-value was calculated by the one-sided paired Student’s t-Test, * *p* < 0.05, ** *p* < 0.01. **c**, The genotype of mutation alleles in T_0_ stable transformation plants. **d**, Base editing frequencies at different protospacer positions by ABE8e at a target site in rice T_0_ lines. **e**, Presence of TA motifs at the target site appears to increase gRNA-dependent off-target A-to-G editing. **f**, The frequency of A-to-G SNV with different di-nucleic acids in T_0_ stable transformation plants.

### Investigation of the somaclonal variation production timeline in rice tissue culture

Since most SNVs (except those from ABE8e-expressing plants) and INDELs are derived from tissue culture, it would be helpful to understand the genesis mechanism and timeline for somaclonal variation. Like many other plants, rice genome editing involves the generation of embryogenic callus, followed by *Agrobacterium* mediated transformation and regeneration[37]. We reasoned that somaclonal variation mutations would be collectively generated before (termed as ‘Phase I somaclonal variation’) and after *Agrobacterium* mediated transformation (termed as ‘Phase II somaclonal variation’) (**Fig. 6a**). Based on the WGS data, we mapped all the T-DNA insertion sites to the rice genome among all the T_0_ lines. Although most plants contained only one T-DNA insertion, 16 plant pairs shared the same T-DNA insertion for each pair (**Fig. 6b**), suggesting each pair of these plants were derived from the same T-DNA transformation event. We hypothesize that shared mutations among such plant pairs would largely represent Phase I somaclonal variations. Our analysis largely confirmed this as the T_0_ plants that share the same T-DNA insertion sites showed high proportion of shared mutations (**Fig. 6c and Supplementary Fig. 11**). Although the numbers of shared mutations for the T_0_ lines with the same T-DNA insertions vary greatly (from 23 to 168), the average number (98) is significantly higher than the average number of shared mutations (7.4) among T_0_ lines with diverse T-DNA insertion sites (**Fig. 6d**).

**Figure 6.**
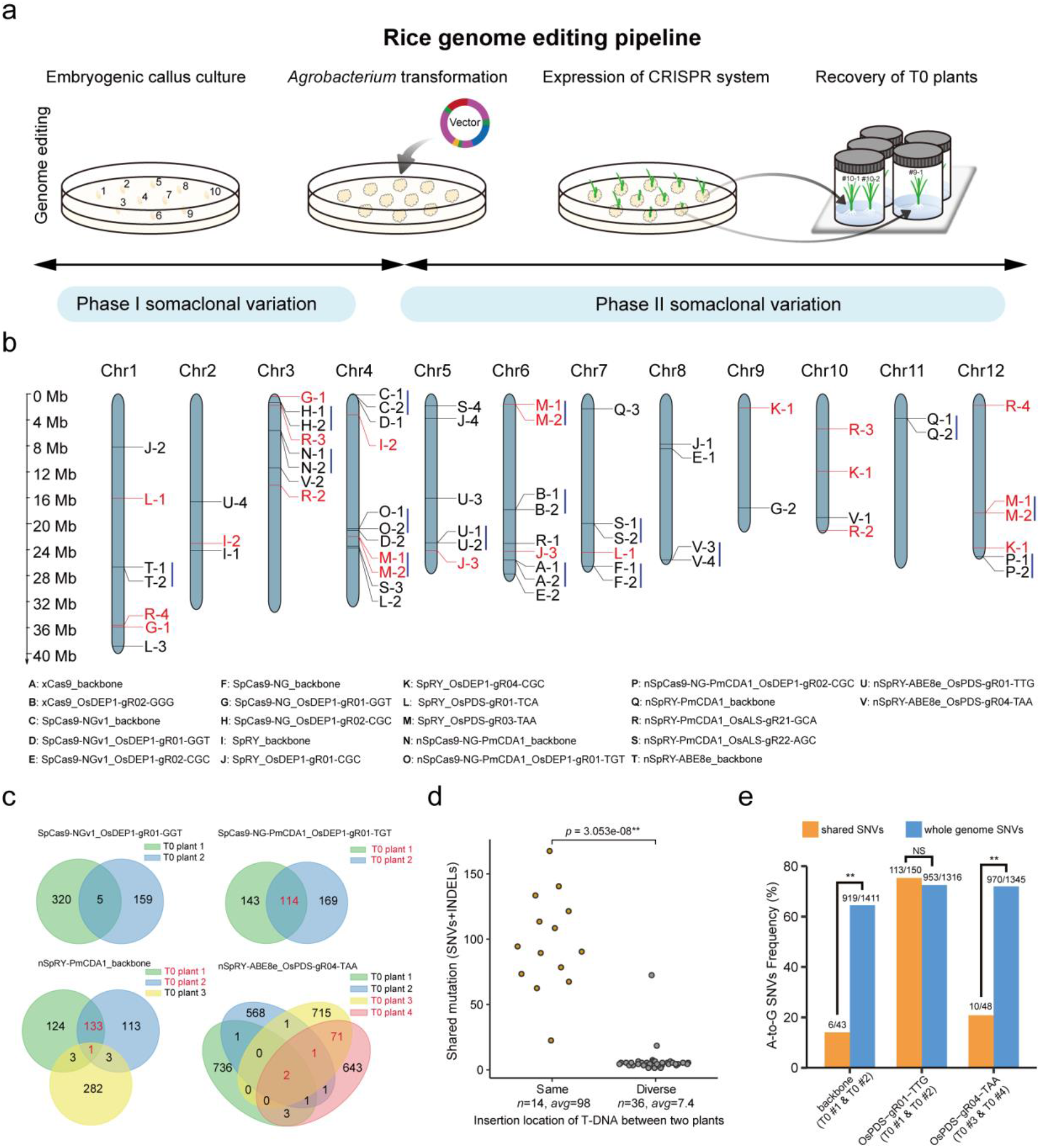
Investigation of somaclonal variation production in rice tissue culture. **a**, A model that divides the generation of somaclonal variation into two phases, which points to potential of minimizing Phase II somaclonal variation with the use of morganic factors to accelerate plant regeneration. **b**, Genome-wide mapping of T-DNA integration sites for all T_0_ lines. Constructs that contain more than one T-DNA integration site are highlighted in red. The two T_0_ lines that carry the same T-DNA integration site were grouped by a solid line on the right, indicating they are from the same transgenic event. **c**, Four examples for the analysis of T_0_ lines for shared mutations revealed by WGS. The T_0_ lines resulting from the same transgenic event (highlighted in red) share a significant portion of mutations (termed Phase I somaclonal variation). **d**, T_0_ lines with the same T-DNA integration sites share an average of 98 mutations, while T_0_ lines with different T-DNA integration sites barely share any mutations. **e**, the frequency of A-to-G SNVs in shared SNVs and whole genome SNVs from the nSpRY-ABE8e T_0_ lines with the same transgenic events, the number above of each bar represents A-to-G SNVs versus all SNVs in a pair of T_0_ lines. *p*-value was calculated by the Wilcoxon rank sum test, * *p* < 0.05, ** *p* < 0.01, NS represents not significant.

We next sought to understand the timeline of genome editing in the context of Phase II somaclonal variation production (**Fig. 6a**). We took advantage of the genome-wide off-target editing by ABE8e and identified three T_0_ plant pairs that were derived from the same transgenic events, based on the shared T-DNA insertion sites (**Fig. 6b**). In all three cases, the sum of whole genome SNVs are more than 1300, with about 70% being A-to-G mutations (**Fig. 6e**), consistent with the genome wide A-to-G off-target mutations by ABE8e (**Fig. 4**). If the ABE8e-based off-target editing were to occur before the transformed callus being developed into two T_0_ lines, the shared mutations between the two T_0_ lines would contain a high percentage of A-to-G mutations. This is indeed the case for the two T_0_ lines edited by nSpRY-ABE8e at OsPDS-gR01-TTG site, where over 70% shared mutations were A-to-G mutations (**Fig. 6e**). For the two remainder cases, about 20% total shared mutations among the two single-event T_0_ lines were A-to-G mutations (**Fig. 6e**), indicating most of the A-to-G off-targeted mutations in these lines were largely independently induced by the same ABE8e transgenic event. These data suggest variable timelines for genome editing to occur in the developmental stage that generates Phase II somaclonal variation. The collective analyses here elucidate the details and timelines of genome editing and somaclonal variation in rice tissue culture: About 100 mutations are Phase I somaclonal variation mutations and about 253 (ranging from 62 to 854) mutations are Phase II somaclonal variation mutations; Genome editing can occur at different timepoints during the Phase II tissue culture stage.

## Discussion

PAM-relaxed Cas9 variants such as SpCas9-NG and SpRY greatly increase the targeting scope in plant genome editing[19–21, 24–26, 28–33]. However, off-target risks also increase with their relaxed PAM restriction and tendency for T-DNA self-editing[19, 35]. Based on WGS analyses in rice, we have found very few off-target mutations induced by SpCas9-NG, SpRY and their derived CBEs based on PmCDA1, a highly specific cytidine deaminase[12]. Our WGS analyses also revealed that SpRY and its derived base editors had higher tendency than SpCas9-NG editors to self-edit their T-DNA[19, 35]. Yet, very limited numbers of off-target mutations were detected in the edited plants by the *de novo* generated new gRNAs. Hence, our results benchmark these genome editing tools for broadened editing scope without significant off-target effects in plants.

The development of the highly processive ABE8e[17, 18] has greatly boosted precise adenine base editing in plants, with up to 100% editing efficiency and extremely low occurrence of INDEL byproducts, which collectively contributed to high frequency of homozygous editing in plants within a single generation[19–23]. Recently, transcriptome-wide analysis in human cells revealed off-target A-to-I conversions caused by ABE8e at the RNA level[18], a phenomenon that was previously reported for ABE7.10[38]. However, significant genome-wide off-target effects have not been previously reported for ABE8e in any organism. Remarkably, we discovered substantial genome-wide off-target effects induced by ABE8e in rice, ~500 A-to-G off-target mutations generated per plant (**Fig. 4a and 4d**). These off-target mutations greatly outweigh the somaclonal variation mutations, presenting a significant implication for the use of ABE8e in plant research. Unlike RNA mutations which are transient and non-inheritable, the resulting A-to-G mutations at the DNA level are largely inheritable (**Supplementary Fig. 9**). Such off-target effects of ABE8e must be addressed before its safe use in plant genetics and crop breeding. Encouragingly, engineered point mutations in the adenosine deaminase have been shown to reduce transcriptome off-target effects by ABE7.10[38], ABE8e[18] and other ABE8 variants[39]. It awaits further testing whether genome-wide off-target A-to-G conversions could be largely mitigated by adopting a highly specific ABE8e variant that carries a promising mutation such as V106W[18, 39].

Interestingly, we found that ABE8e favors editing of TA motifs on DNA, which is consistent with the previous observation that ABE7.10 prefers TA motifs for off-target editing on RNA[38]. Importantly, we found that such a TA motif preference by ABE8e also applies to the target sequence. Hence, this exciting discovery can be applied to improve on-target editing by ABE8e or its further engineered variants by intentionally targeting ‘A’ in a TA motif to achieve high editing efficiency. A CBE was previously used to fine-tune gene expression in strawberry to increase the sugar content[40]. Given the high abundance of TA motifs in the cis regulatory elements (e.g., the TATA box) of many plant genes, ABE8e would be a promising tool for engineering quantitative trait variation by editing cis regulatory elements, an innovative genome editing application that has been conventionally achieved with the Cas9 nuclease(s)[7, 10, 41].

Our WGS analyses, along with the previous studies[11, 12, 16, 34, 42, 43], uncovered the scale of somaclonal variation derived from the tissue culture process, which by itself is a bottleneck for genome editing in plants[44]. Since somaclonal variation is present in all genome-edited plants that are generated by tissue culture, effective strategies are needed to reduce such background mutations, of which many are germline-transmittable[34]. Here, we took a unique approach to investigate the generation of somaclonal variation before and after *Agrobacterium* mediated transformation, which should be applicable to other plants. For the Phase I somaclonal variation mutations, existing before plant transformation (**Fig. 6a**), we may have limited means of reducing them. However, there are often more Phase II somaclonal variation mutations generated, which occur after *Agrobacterium* mediated transformation. We hypothesize that Phase II somaclonal variation may be reduced by accelerating plant regeneration with the expression of morphogenic or growth factors, as recently demonstrated in different plant species[45–47]. It will be promising to test this idea.

In summary, the comprehensive WGS analyses of PAM-relaxed Cas9 nucleases and their derived base editors revealed highly specific genome editing in rice. However, ABE8e, despite its promise for highly efficient and high-purity base editing, showed substantial genome-wide off-target A-to-G conversions that are independent of gRNAs. This study also points to promising approaches of enhancing on-target and reducing off-target A-to-G editing by ABE8e or its variants, as well as potentially reducing Phase II somaclonal variation in genome-edited plants.

## Methods

### Plant material and growth condition

The Nipponbare rice cultivar (*Oryza sativa* L. ssp. Japonica cv. Nipponbare) was used in this study as the WT control and transformation host. All plants for the WGS assay were grown in growth chambers under a controlled environmental condition of 60% relative humidity with a 16/8 h and 32/28 °C regime for under the light/dark cycle.

### Construction of T-DNA vectors

The PAM-relaxed CRISPR-Cas9 plant genome editing systems used in this study were reported in our previous studies [19, 25]. Target sites were inserted by Golden Gate reaction using BsaI HF v02 and T4 DNA Ligase (New England Biolabs) per our previous description[48–50]. Briefly, the synthesized pair oligos (10μM) were annealed and cool down to room temperature (23 °C). The annealed mixture was diluted to 50 nM for a total 15 cycles in the Golden Gate reaction[49, 50]. The reaction mixture was transformed to *Escherichia coli* DH5α competent cells followed by miniprep and Sanger sequencing.

### Rice transient and stable transformation

Rice protoplast isolation, transformation and editing activity evaluation were performed as described previously[51–53]. The *Agrobacterium* mediated rice stable transformation was based on previously published protocols with minor modifications[54–56]. Briefly, the rice calli was induced and the binary T-DNA vectors were transformed into *Agrobacterium tumefaciens* EHA105 strain. The transformed EHA105 strain was cultured in the flask until the OD600=0.1 at 28 °C and collected by centrifuge. The collected *Agrobacterium* was resuspended with AAM-AS medium for calli transformation. After 3 days of co-incubation, the transformed calli were washed by sterile water and transferred to N6-S solid medium for 14 days under continuous light at 32 °C. The grown calli were collected and incubate at REIII solid medium. After a 14-day regeneration, the newly grown individual plants were transferred to HF solid medium for root induction. Then, the generated plants were moved into pods and grown in soil at growth chamber under 18 h light at 32 °C and 6 h dark at 28 °C. After 4 weeks’ growth, the leaf was collected both for targeted mutagenesis assay and whole genome sequencing.

### Mutation detection and analysis

The genomic DNA was extracted using the CTAB method[57]. About 100 ng genomic DNA and a 50 uL PCR reaction was used to amplify the transgene and target sequence for detection of transgenic plants and genome editing events. The oligos used in this study were shown in **Supplementary Table 4**. PCR was done with 2xRapid Taq Mix (Vazyme) and examined using SSCP strategy[58]. The genotype at the target sites of each plant was confirmed by Sanger sequencing.

### Whole genome sequencing and data analysis

One gram of fresh leaves was obtained from each edited rice plant for WGS. Genomic DNA was extracted using Plant Genome DNA Kit (Tiangen). All plant samples were sequenced by the Illumina NovaSeq platform (Novogene, Beijing, China). The average sequencing clean data generated for each sample was 20 Gb, with the average depth being ~50X to 70X. For data processing, adapters and low quality reads were first trimmed and filtered using SKEWER (v. 0.2.2)[59]. Cleaned reads were then mapped to rice reference sequence TIGR7 (MSU7) with BWA mem (v. 0.7.17) software[60]. Picard (https://broadinstitute.github.io/picard/) software (v. 2.22.4) and Samtools (v. 1.9)[61] were employed to mark duplicate reads and generate sort BAM files, respectively. The Genome Analysis Toolkit (GATK v. 3.8)[62] was applied to realign the reads near INDELs and recalibrate base quality scores against known SNPs and INDELs databases (http://snp-seek.irri.org/). After the raw BAM files were processed by GATK, analysis-ready BAM files were generated. To identify genome-wide somatic mutations with high confidence, we applied three software each to identify SNVs and INDELs, respectively. Whole genome SNVs were detected by LoFreq (v. 2.1.2)[63], MuTect2[64] and VarScan2 (v. 2.4.3)[65]. Whole genome INDELs were detected by MuTect2[64], VarScan2 (v. 2.4.3)[65] and Pindel (v. 0.2)[66]. The Bedtools (v. 2.27.1)[67] was used to obtain overlapping SNVs/INDELs among replicates or different software. SNVs and INDELs identified by all three corresponding software were retained for the further analysis. Cas-OFFinder *in silico* (v. 2.4)[36] was used to predicted putative off-target sites in the rice genome. The PAM type of SpRY, SpCas9-NG and xCas9 were set to NNN, NGN and NGN, respectively, allowing up to 5-nt mismatches in the protospacer. IGV (v. 2.8.4) software[68] was applied to visualize discovered mutations with the generated BAM and VCF files. To identify the insertion locations of T-DNA in each line, the cleaned reads were first aligned to the rice reference genome and vector sequences simultaneously. Then, the BAM files were visualized using the IGV software and ‘Group Alignments by’ mode was set to ‘chromosome of mate’ in IGV. Lastly, each T-DNA insertion site was confirmed by manual checking of paired reads aligned to both vector sequences and specific chromosomes. The genome-wide distribution of mutations was drawn by Circos (v 0.69)[69]. The adjacent 3-bp sequences of the A-to-G SNVs were extracted from the reference genome sequence, and then submitted to WebLogo3 (http://weblogo.threeplusone.com/)[70] to plot motif weblogo. Data processing, analyses, and figure plotting were completed by using R and Python.

### Data availability

The WGS data have been deposited in the Sequence Read Archive in National Center for Biotechnology Information (NCBI) under the accession number PRJNA792795 and Beijing Institute of Genomics Data Center (http://bigd.big.ac.cn) under BioProject PRJCA007564.

## Acknowledgements

This research was supported by the Sichuan Science and Technology Program (award no. 2021JDRC0032, 2021YFH0084 and 2021YFYZ0016) to J.Z. and Y.Z., the National Natural Science Foundation of China (award no. 32101205, 32072045 and 31960423) to X.T. and X.Z., the Open Foundation of Jiangsu Key Laboratory of Crop Genetics and Physiology (award no. YCSL202009) to J.Z, Y.Z and T.Z. It is also supported by the National Science Foundation Plant Genome Research Program (award no. IOS-2029889) and the U.S. Department of Agriculture Biotechnology Risk Assessment Grant Program competitive grant (award no. 2018-33522-28789) to Y.Q. S.S. is a Foundation for Food and Agriculture Research Fellow.

## Author contributions

Y.Z., T.Z. and Y.Q. conceived and designed the experiments. Q.R., Z.Z., X.T. and S.S. made the vectors for rice transformation. Q.R. and Z.Z. conducted rice protoplast transformation and data analysis. Q.R., Z.Z., L.L., S.X. and X.Z. did the rice stable transformation and mutagenesis assays. Q.R., Z.Z., L.L., S.X. and J.Z. prepared rice seedling samples for WGS. Y. W. and G. L. performed WGS data analysis and generated the figures. Y. H., Y. B. and S. L. assisted with data analysis. Y.Z., T.Z. and Y.Q. supervised the research and wrote the manuscript. All authors participated in discussion and revision of the manuscript.

## Additional information

Supplementary information is available for this paper.

---

**Supplementary Fig 1.**
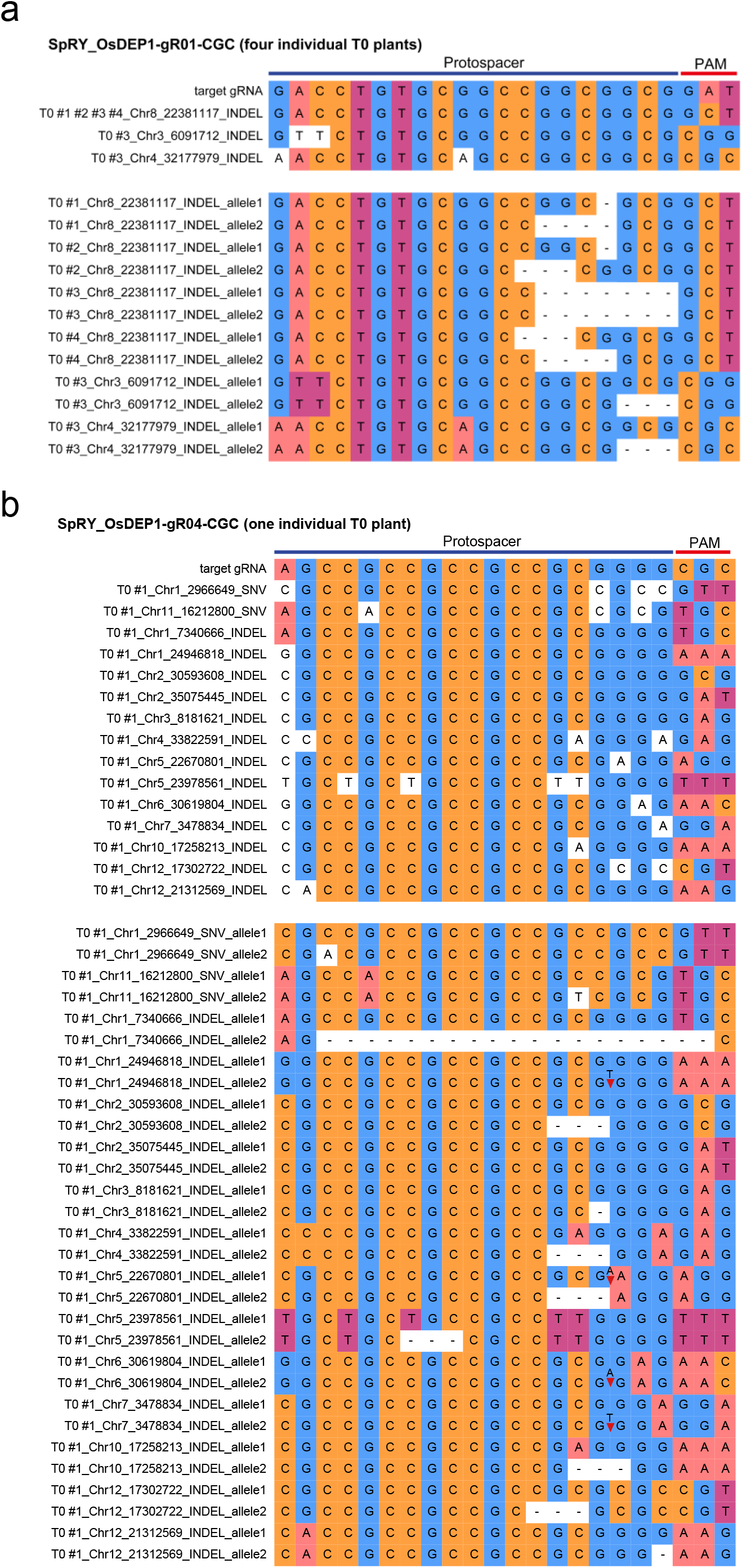
Guide RNA-dependent off-target mutagenesis by SpRY. **a-b**, gRNA-dependent off-target mutations in edited T0 lines at the OsDEP1-gR01-CGC site **(a)** and OsDEP1-gR04-CGC site **(b)**.

**Supplementary Fig 2.**
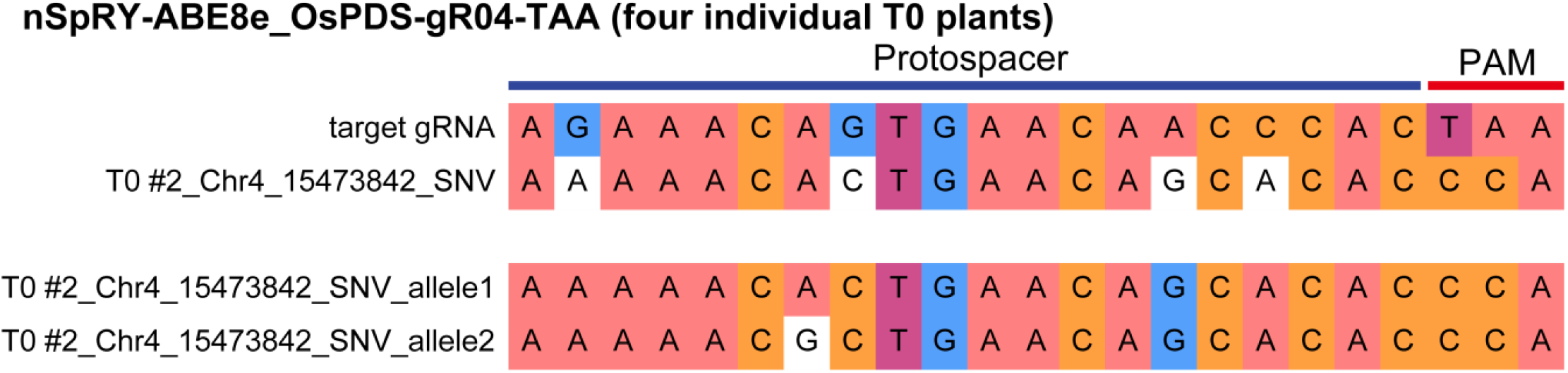
Guide RNA-dependent off-target mutagenesis by nSpRY-ABE8e at OsPDS-gR04-TAA site.

**Supplementary Fig 3.**
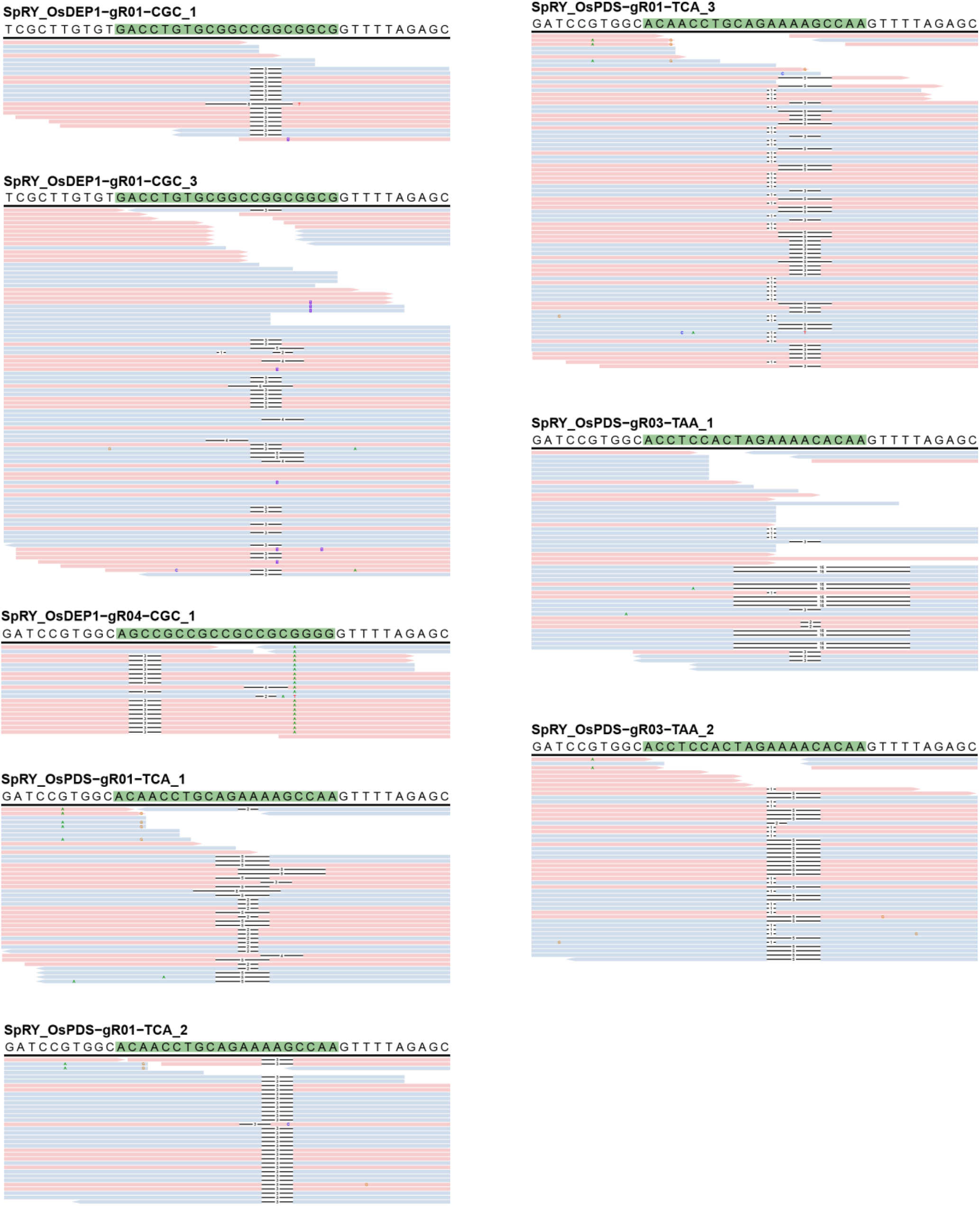
Sequencing reads indicative for T-DNA self-editing by SpRY constructs. Protospacer sequences are marked by green rectangles. Insertions are marked by purple boxes. Deletions are marked by black dashes. Mismatches are marked by colored bases.

**Supplementary Fig 4.**
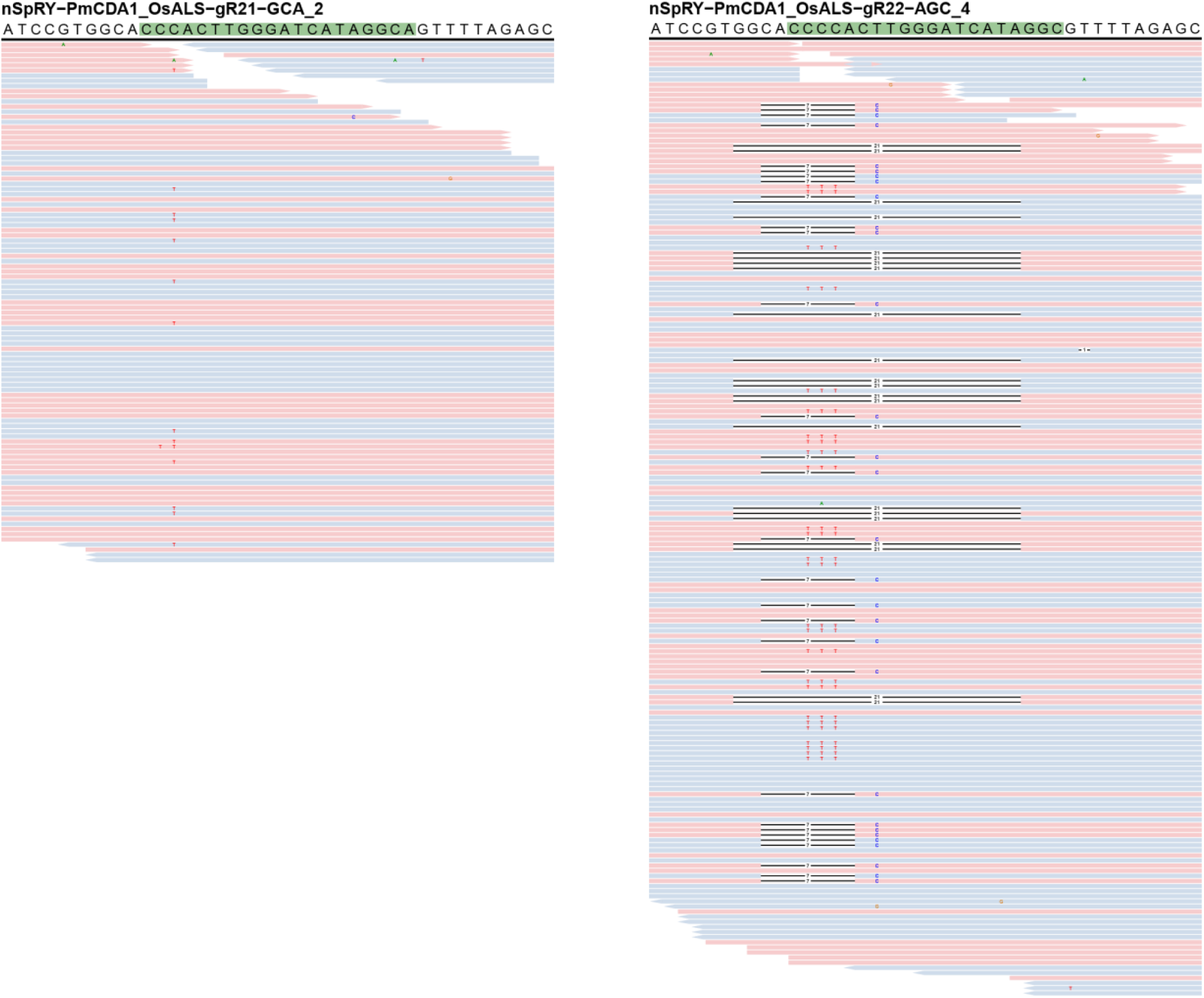
Sequencing reads indicative for T-DNA self-editing by nSpRY-PmCDA1 constructs. Protospacer sequences are marked by green rectangles. Insertions are marked by purple boxes. Deletions are marked by black dashes. Mismatches are marked by colored bases.

**Supplementary Fig 5.**
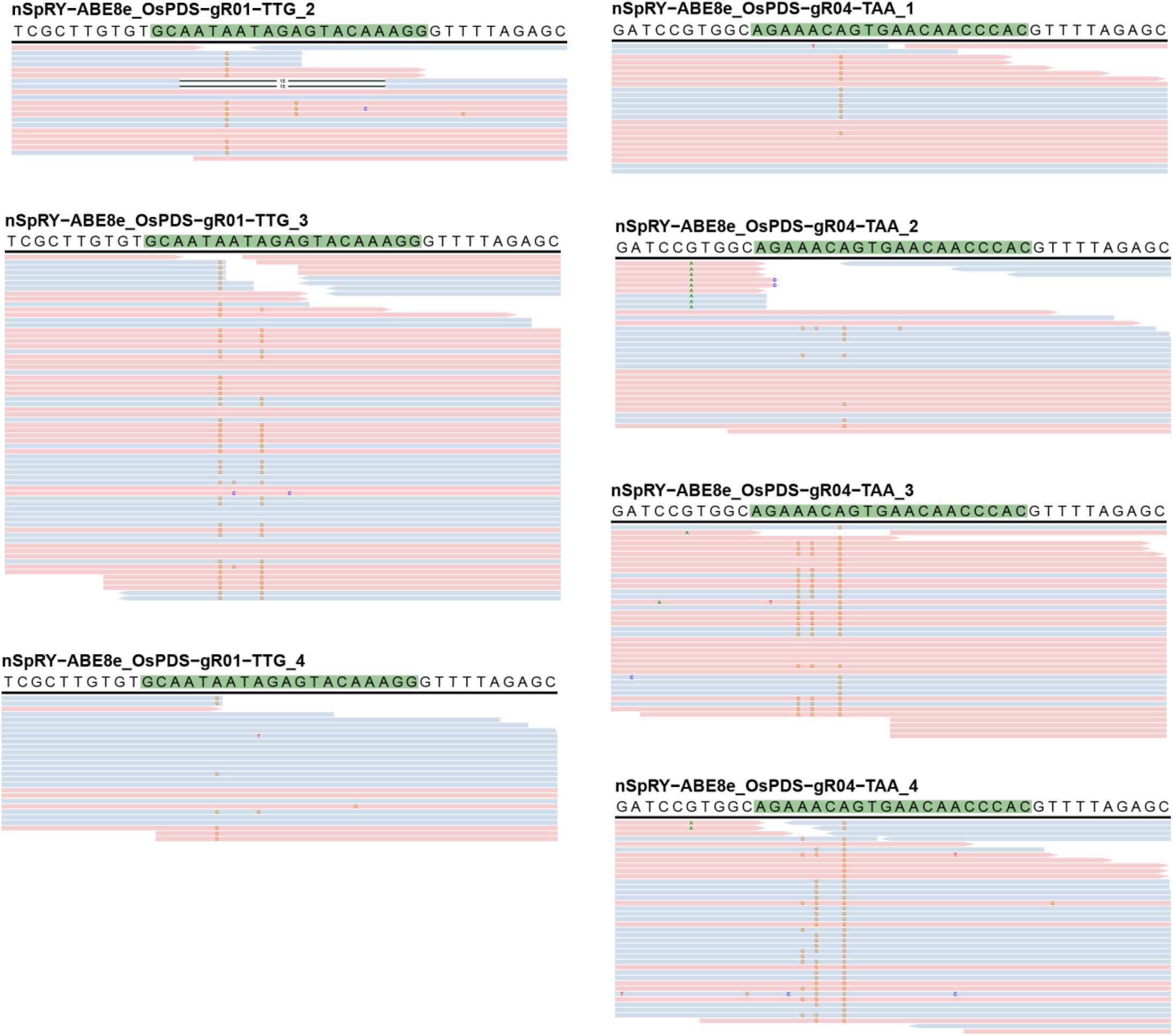
Sequencing reads indicative for T-DNA self-editing by nSpRY-ABE8e constructs. Protospacer sequences are marked by green rectangles. Insertions are marked by purple boxes. Deletions are marked by black dashes. Mismatches are marked by colored bases.

**Supplementary Fig 6.**
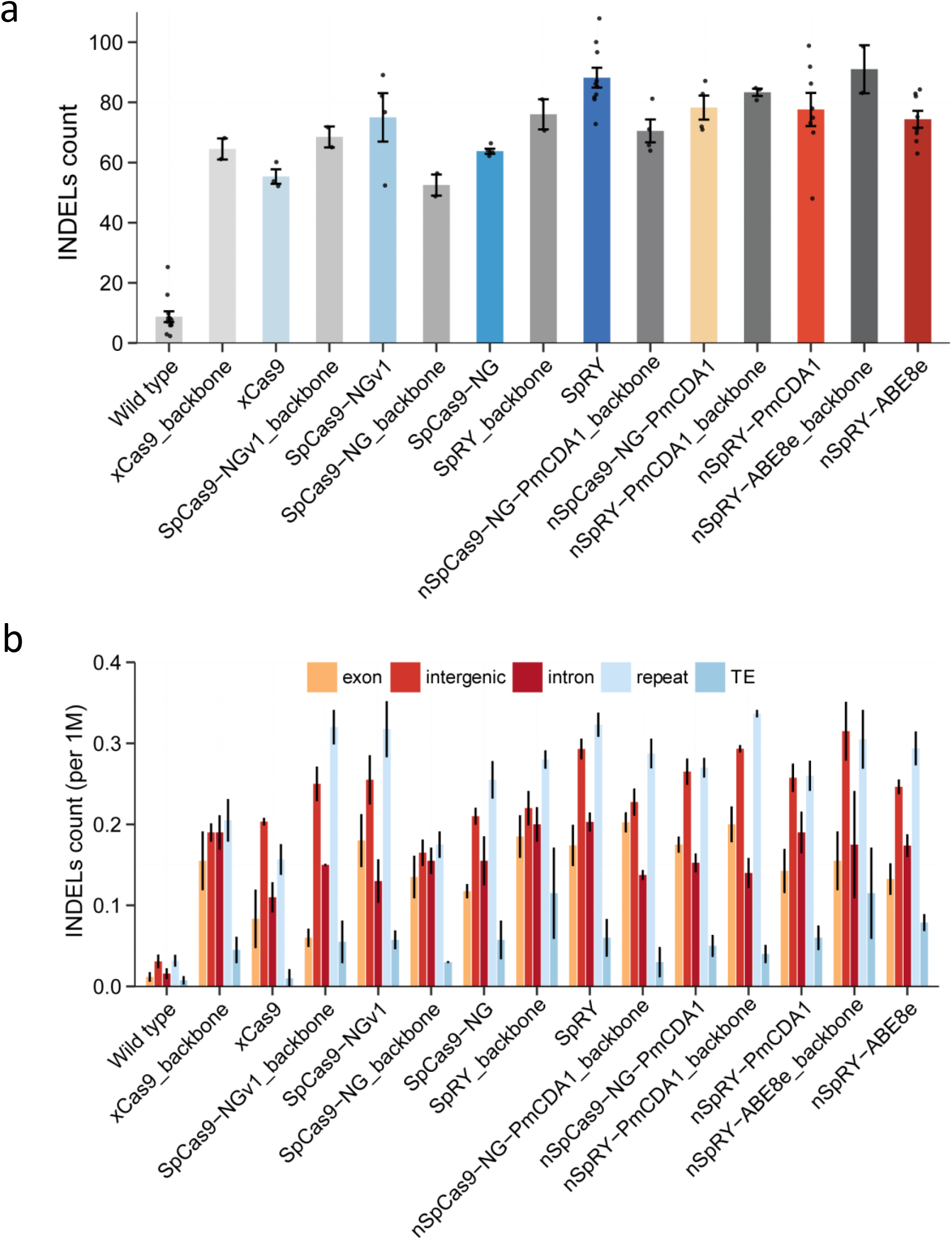
INDEL mutations in all sequenced samples and their genome-wide distributions. **a**, Number of INDELs identified in all 58 sequenced samples. **b**, Average number of SNV mutations per 1 Mbp genomic region. Error bars represent s.e.m and the dots represent individual plants.

**Supplementary Fig 7.**
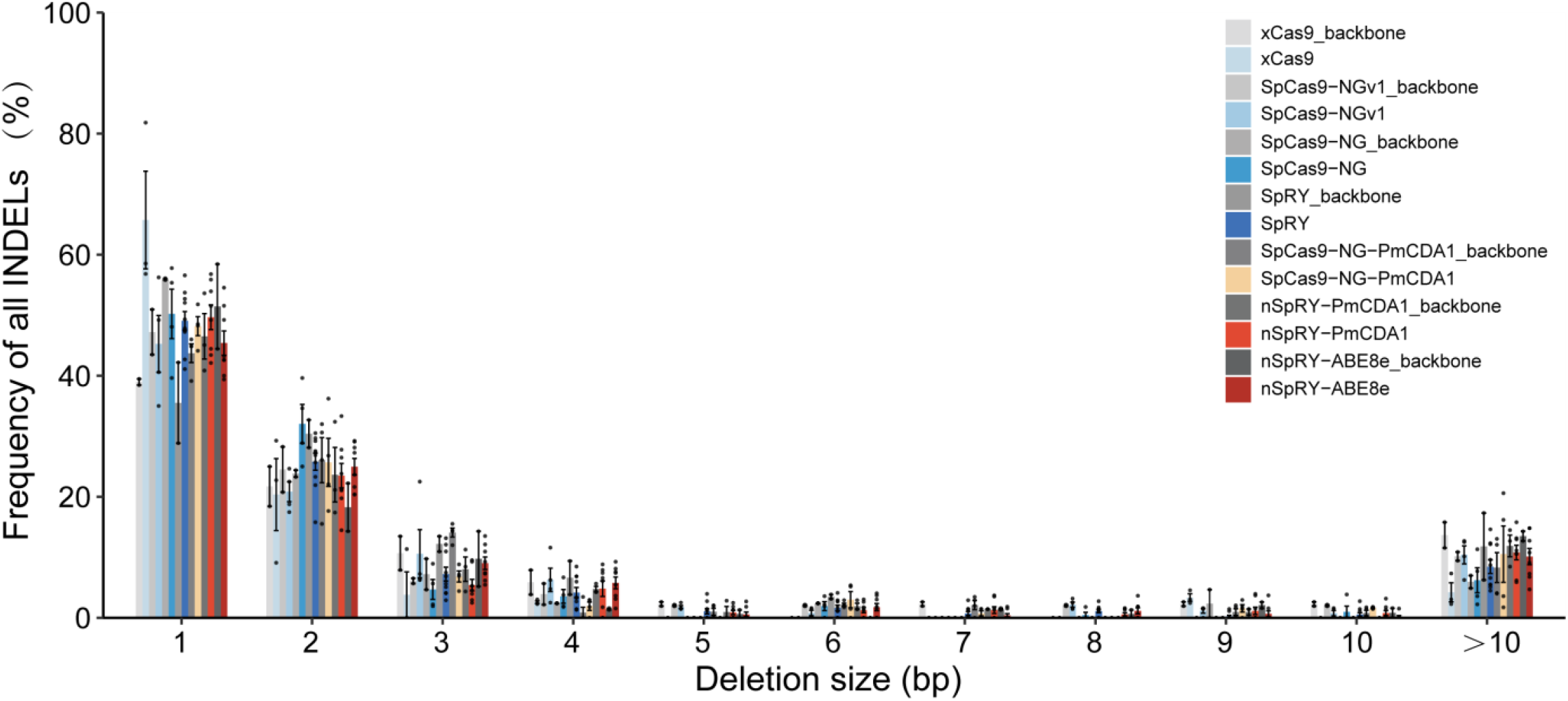
Comparison of deletion sizes among all mutations induced by different genome editing systems. Error bars represent s.e.m and the dots represent individual T_0_ plants.

**Supplementary Fig 8.**
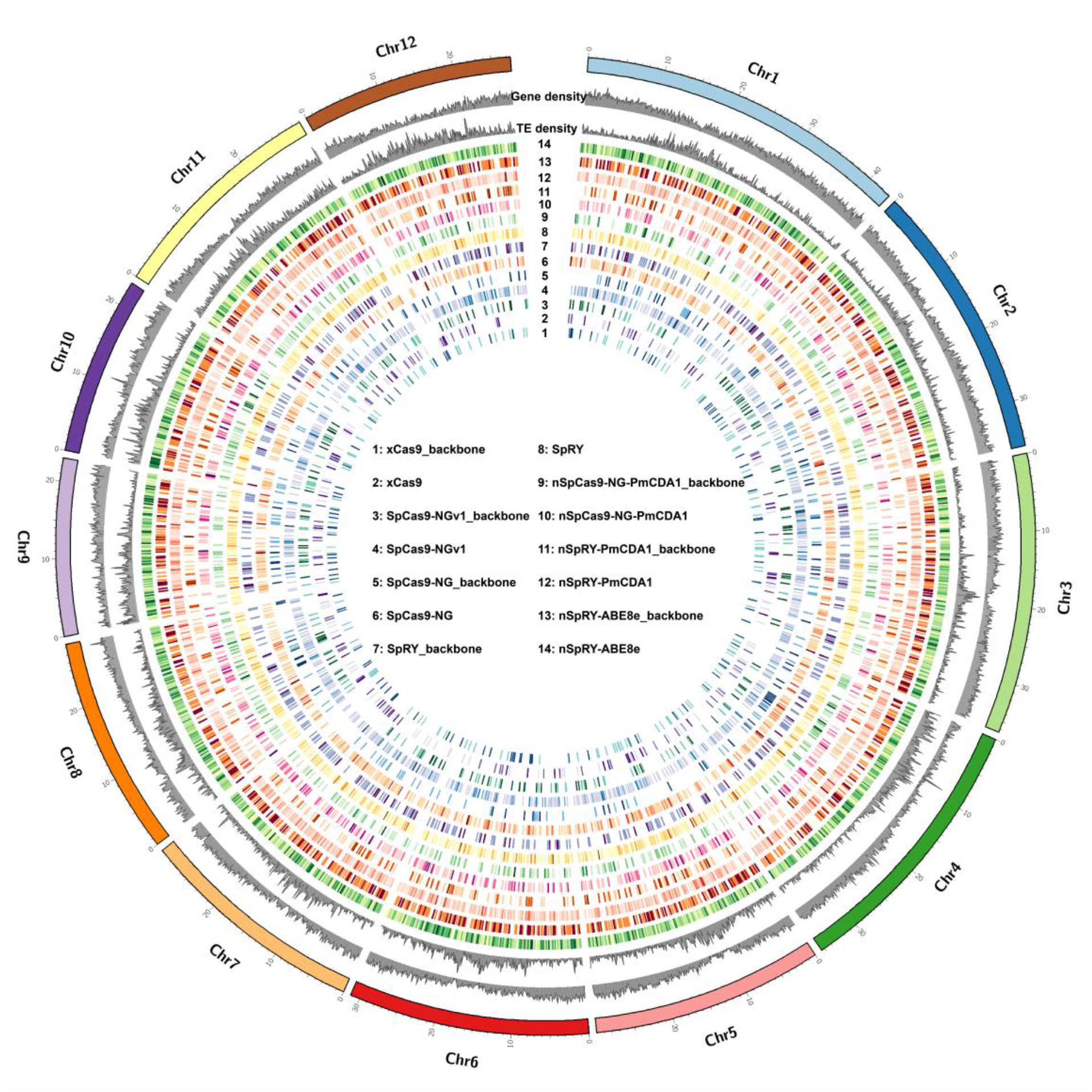
Genome-wide distribution of mutations (SNVs+INDELs) from all sequenced sample.

**Supplementary Fig 9.**
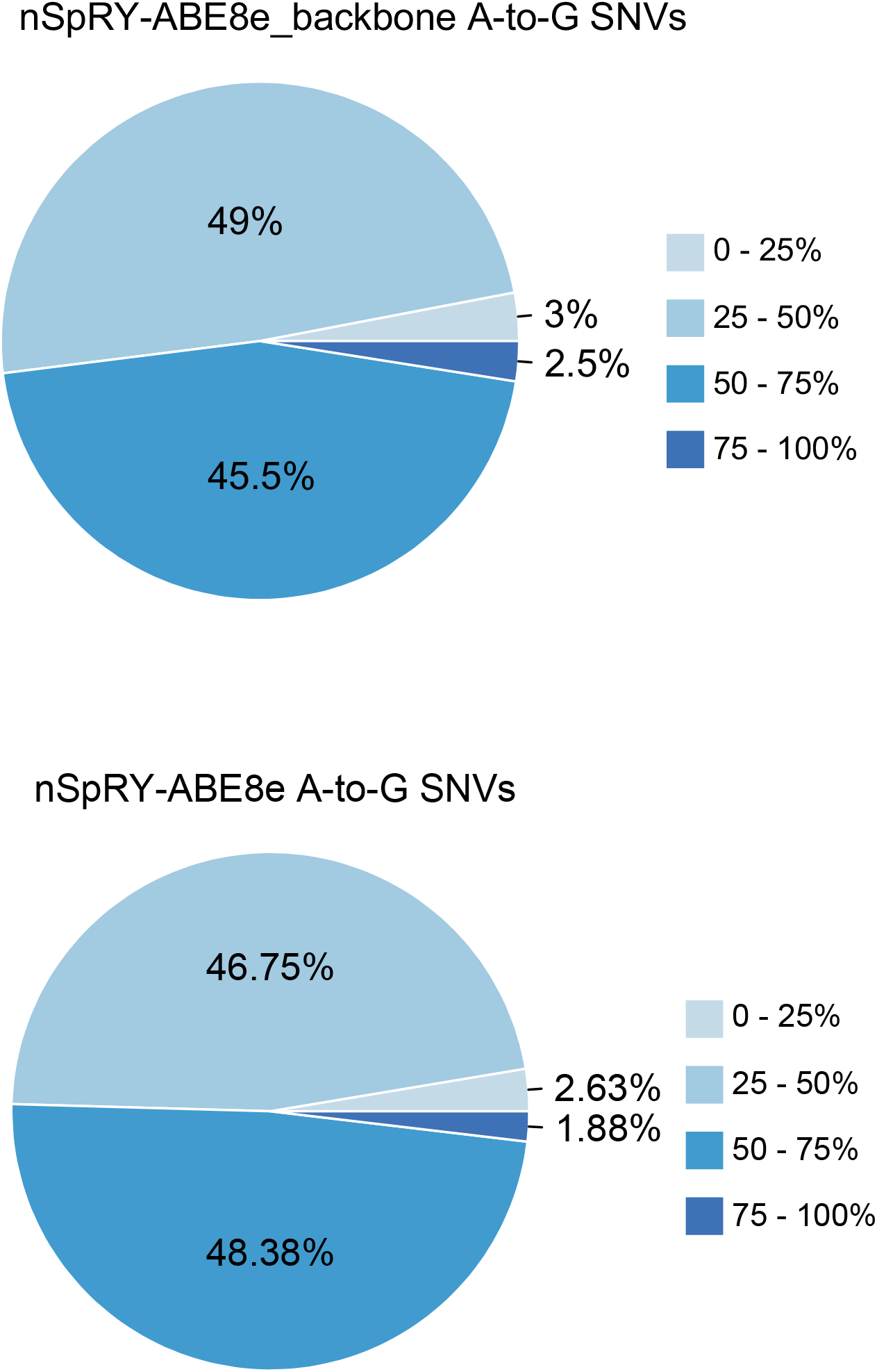
Allele frequencies of A-to-G SNVs identified in nSpRY-ABE8e (*n*=8) and nSpRY-ABE8e_backbone (*n*=2) samples.

**Supplementary Fig 10.**
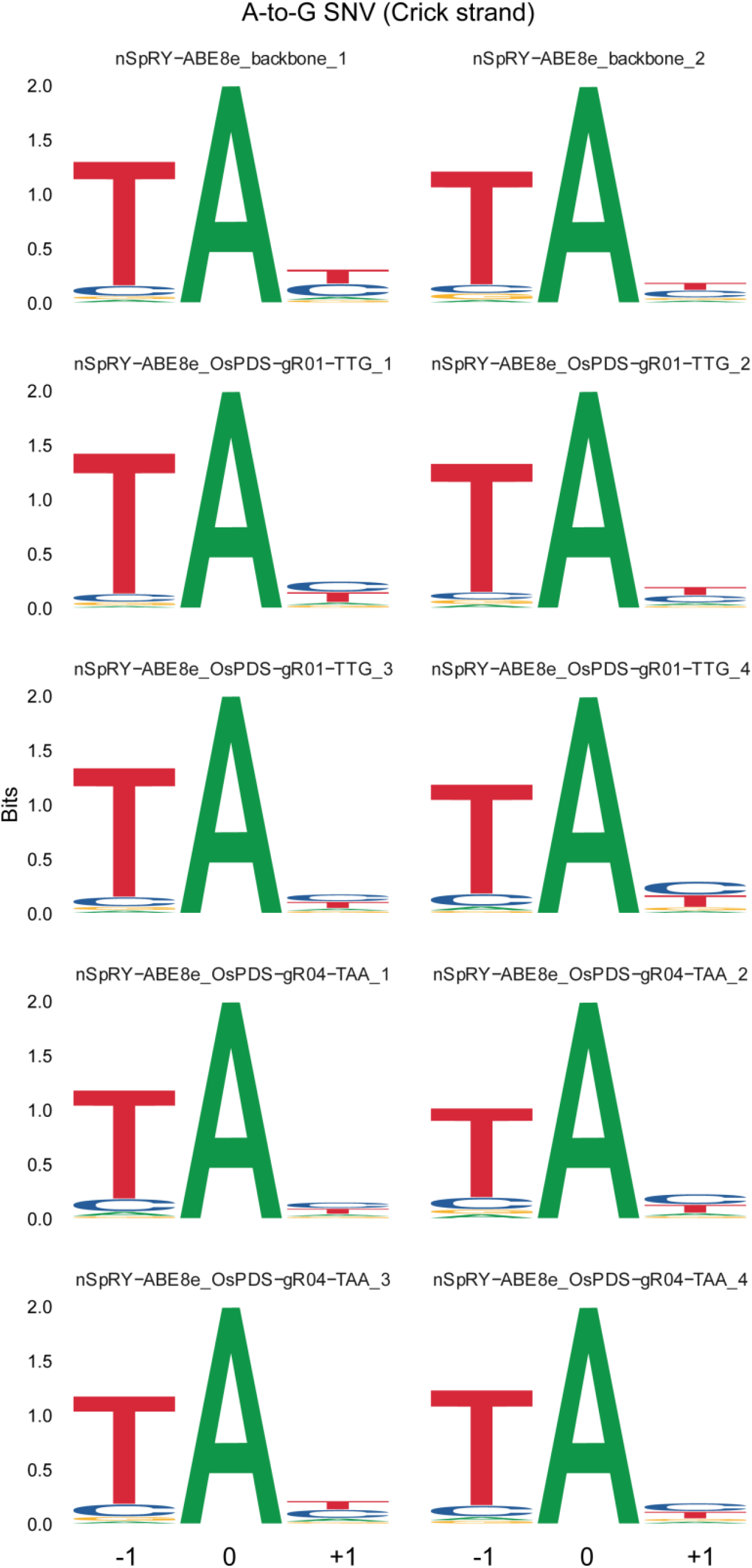
Sequence signature of ABE8e based genome-wide off-target mutations. Preference of a TA motif by ABE8e at gRNA-independent off-target A-to-G base editing in Crick strand. The ‘0’ indicates the A-to-G conversion position.

**Supplementary Fig 11.**
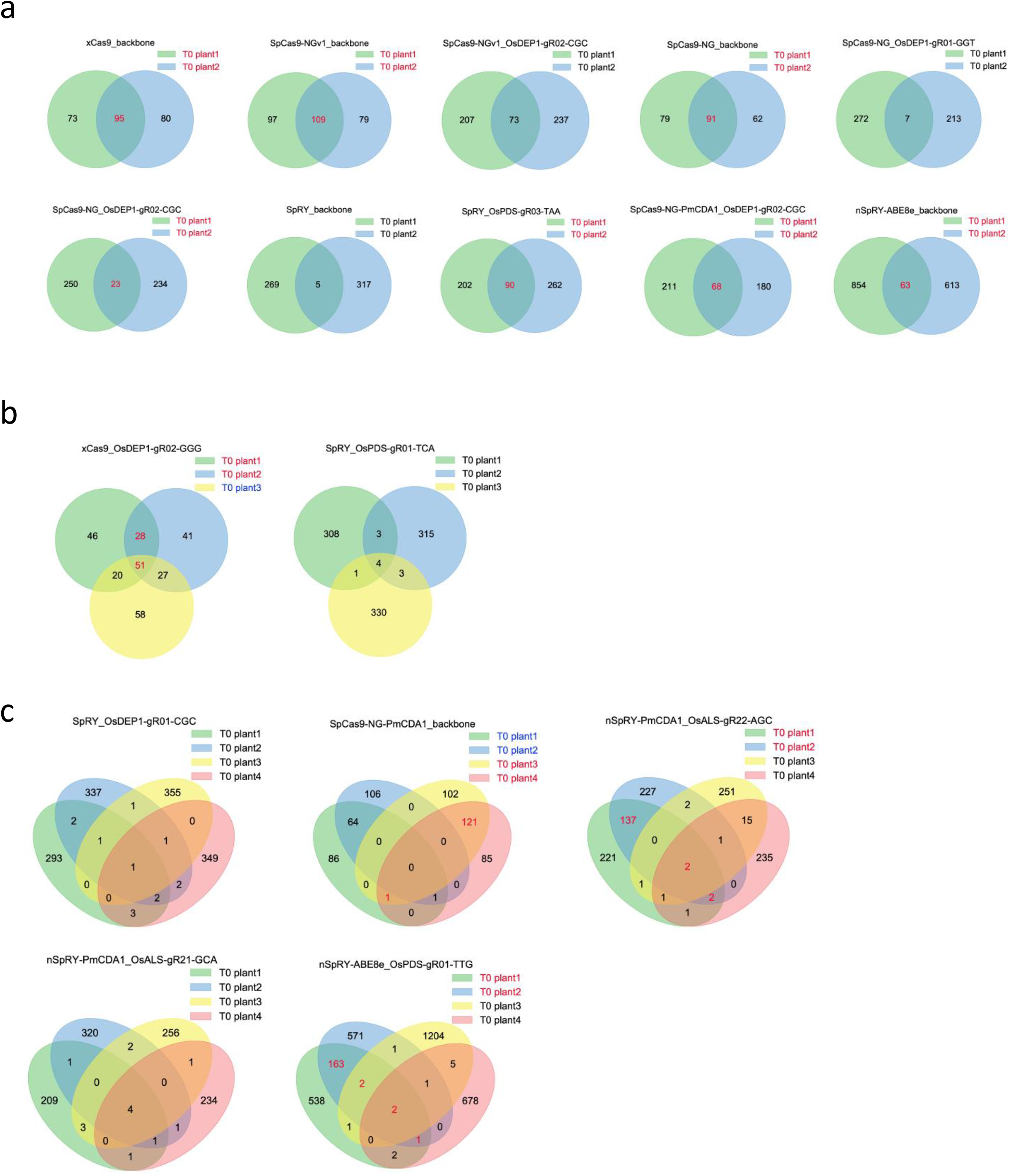
Venn diagram showing mutations shared between individual plants. Each circle or oval represents an individual T_0_ plant. Constructs that resulted in two T_0_ lines (**a**), three T_0_ lines (**b**), and four T_0_ lines (**c**) were shown. The T_0_ lines resulting from the same transgenic event are marked in red.

